# A needle in a haystack: a new metabarcoding approach to survey diversity at the species level of Arcellinida (Amoebozoa: Tubulinea)

**DOI:** 10.1101/2022.07.12.499778

**Authors:** Rubén González-Miguéns, Emilio Cano, Antonio Guillén-Oterino, Antonio Quesada, Daniel J.G. Lahr, Daniel Tenorio-Rodríguez, David de Salvador-Velasco, David Velázquez, María Isabel Carrasco-Braganza, R. Timothy Patterson, Enrique Lara, David Singer

**Affiliations:** Department of Mycology, Real Jardín Botánico (RJB-CSIC), C/ Moyano 1, 28014 Madrid, Spain; Research Support Unit, Real Jardín Botánico (CSIC), C/ Moyano 1, 28014 Madrid, Spain; Estación Biológica Internacional Duero-Douro (EUROPARQUES-EBI), Buque hidrográfico Helios-Cousteau en el Lago de Sanabria, Castilla y León, Spain; Department of Biology, Universidad Autónoma de Madrid. 28049, Madrid, Spain; Department of Zoology, Institute of Biosciences, University of São Paulo, Brazil; Ottawa-Carleton Geoscience Centre and Department of Earth Sciences, Carleton University, 1125 Colonel By Drive, Ottawa, Ontario, K1S 5B6, Canada; UMR CNRS 6112 LPG-BIAF, Laboratory of Planetology and Geosciences, Angers University and Nantes Université, 2 Boulevard Lavoisier, 49045 Angers, France

**Keywords:** Arcellinida, mitochondrial genes, eDNA, metabarcoding, species-level, testate amoeba

## Abstract

Environmental DNA-based diversity studies have increased in popularity with the development of high throughput sequencing technologies. This permits the potential simultaneous retrieval of vast amounts of molecular data from many different organisms and species, thus contributing to a wide range of biological disciplines. Environmental DNA protocols designed for protists often focused on the highly conserved small subunit of the ribosome gene, that does not permit species-level assignments. On the other hand, eDNA protocols aiming at species-level assignments allow a fine level ecological resolution and reproducible results. These protocols are currently applied to organisms living in marine and shallow lotic freshwater ecosystems, often in a bioindication purpose. Therefore, in this study, we present a species-level eDNA protocol, designed to explore diversity of Arcellinida (Amoebozoa: Tubulinea) testate amoebae taxa, that is based on mitochondrial cytochrome oxidase subunit I (COI). These organisms are widespread in lentic water bodies and soil ecosystems. We applied this protocol to 42 samples from peatlands, estuaries and soil environments, recovering all the infraorders in Glutinoconcha (with COI data), except for Hyalospheniformes. Our results revealed an unsuspected diversity in morphologically homogeneous groups such as Cylindrothecina, Excentrostoma or Sphaerothecina. With this protocol we expect to revolutionize the design of modern distributional Arcellinida surveys. Our approach involve a rapid and cost effective analysis of testate amoeba diversity living in contrasted ecosystems. Therefore, Arcellinida clade have the potential to be established as a model group for an array of theoretical and applied studies.

## 1 Introduction

The use of environmental DNA (eDNA) has revolutionized biodiversity studies (Taberlet et al., 2012; Yoccoz, 2012), culminating with the development of high throughput sequencing (HTS) technologies (Shokralla et al., 2012). The application of HTS to amplicon sequencing (i.e., metabarcoding) allowed the retrieval of massive amount of genetic data from many different organisms. This breakthrough has contributed significantly to academic research, especially in the fields of ecology, evolution or biogeography (Beng & Corlett, 2020; Boenigk et al., 2018; Gaither et al., 2022) and, in applied research, to environmental quality assessment (Lezcano et al., 2017; Pawlowski et al., 2021), amongst others. Nevertheless, to draw sound ecological or biogeographical conclusions based on metabarcoding data, an accurate reference taxonomic database is required. This database must be as exhaustive as possible, with representatives of the main clades that constitute the surveyed groups. This is the case for the most popular eukaryotic metabarcoding marker, the gene coding for the small subunit of the ribosome known as 18S rRNA gene (Berney et al., 2017; del Campo et al., 2018). The eukaryotic 18S reference database PR^2^ (Guillou et al., 2013) is very large and currently comprises (last accessed 06.06.2022) of almost 200,000 taxonomic annotated sequences, which makes possible a relatively accurate placement of newly obtained environmental sequences into known clades. This extensive taxonomic database and the universal presence of 18S rRNA in all eukaryotes has permitted the development of generic eDNA PCR protocols to retrieve a large part of environmental biodiversity without requiring prior knowledge of the taxonomic composition of communities (but see Vaulot et al., 2022). However, despite these attributes the 18S rRNA gene is too conserved to be used for species identification in both micro- and macroscopic sized eukaryotes (Lara et al., 2022; Tang et al., 2012). Due to this the taxonomic resolution of the 18S rRNA gene is usually treated as Operational Taxonomic Units (OTUs), which clusters the sequences with regards to their similarity, but is lumping organisms with divergent ecological functions (Lara et al., 2022). As species are the only truly biologically and ecologically meaningful (and comparable) units of diversity in eukaryotes (Bickford et al., 2007; de Queiroz, 2005; Wiley, 1978), specific protocols designed to discriminate species-level and curated databases are necessary in eDNA biodiversity studies.

Species-level metabarcoding protocols have already been successfully (and routinely) applied in macro-organism (e.g. plants, animals, fungi; Ficetola et al., 2008; Richards et al., 2012). Metabarcoding protocol applications to bioindication (Czechowski et al., 2020), conservation biology (Cilleros et al., 2019; Pfleger et al., 2016), invasive species monitoring (Ardura et al., 2015) or biogeography assessment (West et al., 2021) allow the retrieval of large amounts of molecular data without direct observation of the species, facilitating fieldwork (Christianson et al., 2022). In protists (i.e. the major part of eukaryotic diversity) these protocols are only seldom applied, which relegates related studies to more theoretical-generalist topics (Pawlowski et al., 2016). Still, protists could be subject to applied research, as for instance bioindication of environmental disturbance. These microorganisms are considered excellent bioindicators due to their ubiquity, rapid response to environmental changes and their short lifecycles (Payne, 2013). An important requirement though is that the application of species-level metabarcoding protocols in eDNA studies requires the use of fast evolving molecular markers.

As a starting point to develop species-level metabarcoding, barcoding protocols that allow species level discrimination have been proposed for protists, notably based on the nuclear internal transcribed spacer (ITS), or the mitochondrial cytochrome oxidase subunit I (COI). These former barcodes were first tested against isolated and already characterized organisms, proving that they provide a good resolution at the species level (Heger, Pawlowski, et al., 2011; Kostka et al., 2017; Macher et al., 2021; Moniz & Kaczmarska, 2009). Despite the proven success of these approaches for (organism) barcoding, protocols have been seldom transferred to environmental studies. The exceptions have been ecological research involving diatom barcodes (Kermarrec et al., 2013) and most recently foraminifera (Girard et al., 2022). Both of these groups are excellent ecological and paleoecological bioindicators in marine environments. In addition, diatoms are also studied in freshwater systems, including streams and a variety of shallow lotic settings. Diatoms are also widely used as important proxies for ecological status assessment (e.g., the European Water Framework Directive; WFD; European Commission, 2000). The last decade has seen the development of cutting-edge metabarcoding protocols for diatom profiling that promise to eventually provide rapid, affordable and reliable tools for routine environmental ecological assessment in both freshwater and marine ecosystems, (Kermarrec et al., 2014; Mortágua et al., 2019). Routine and wide-scale application of the technology has not occurred yet though as standardization of eDNA and bioinformatics protocols are still pending (Bailet et al., 2020). Other important groups have barely been studied though with species-level metacobarcoding protocols. We propose here that Arcellinida (Amoebozoa: Tubulinea), also known as lobose testate amoebae, an important intermediary trophic group widespread in lentic water bodies and soil ecosystems, are an ideal test bed upon which to developed as species-level eDNA bioindicators in these environments.

Arcellinida is a group of amoeboid protists with a worldwide distribution, and can be found in soil, freshwater sediments, plankton and, to a lesser extent, marine environments where they inhabit interstitial zones (Smith et al., 2009). These organisms have a characteristic self-constructed test or shell whose size, shape and composition is used for taxonomic identification (Fig. 1). In most studies these organisms are primarily identified and enumerated morphologically and are routinely used to provide insights into both substrate character and the general ecological health of lakes (Patterson & Kumar, 2002). They are routinely used for specific applications such as monitoring perturbations in lacustrine systems including; heavy metal contamination (Nasser et al., 2020; Yang et al., 2011), road salt loading (Cockburn et al., 2020) and eutrophication (Macumber et al., 2020). In terrestrial habitats, they are used in peatland restoration management (Carballeira & Pontevedra-Pombal, 2021; Marcisz et al., 2016; Valentine et al., 2013), or to evaluate the impact of open-cast coal mining on soil (Wanner & Dunger, 2001). There is thus an extensive scientific literature on the sensitivity of these organisms to different environmental perturbations (Freitas et al., 2022). In addition to their use as bioindicator, Arcellinida have also been proposed as a model group for studying microbial biogeography (Heger, Lara, et al., 2011; Singer et al., 2019; Smith & Wilkinson, 2007).

**Figure 1.**
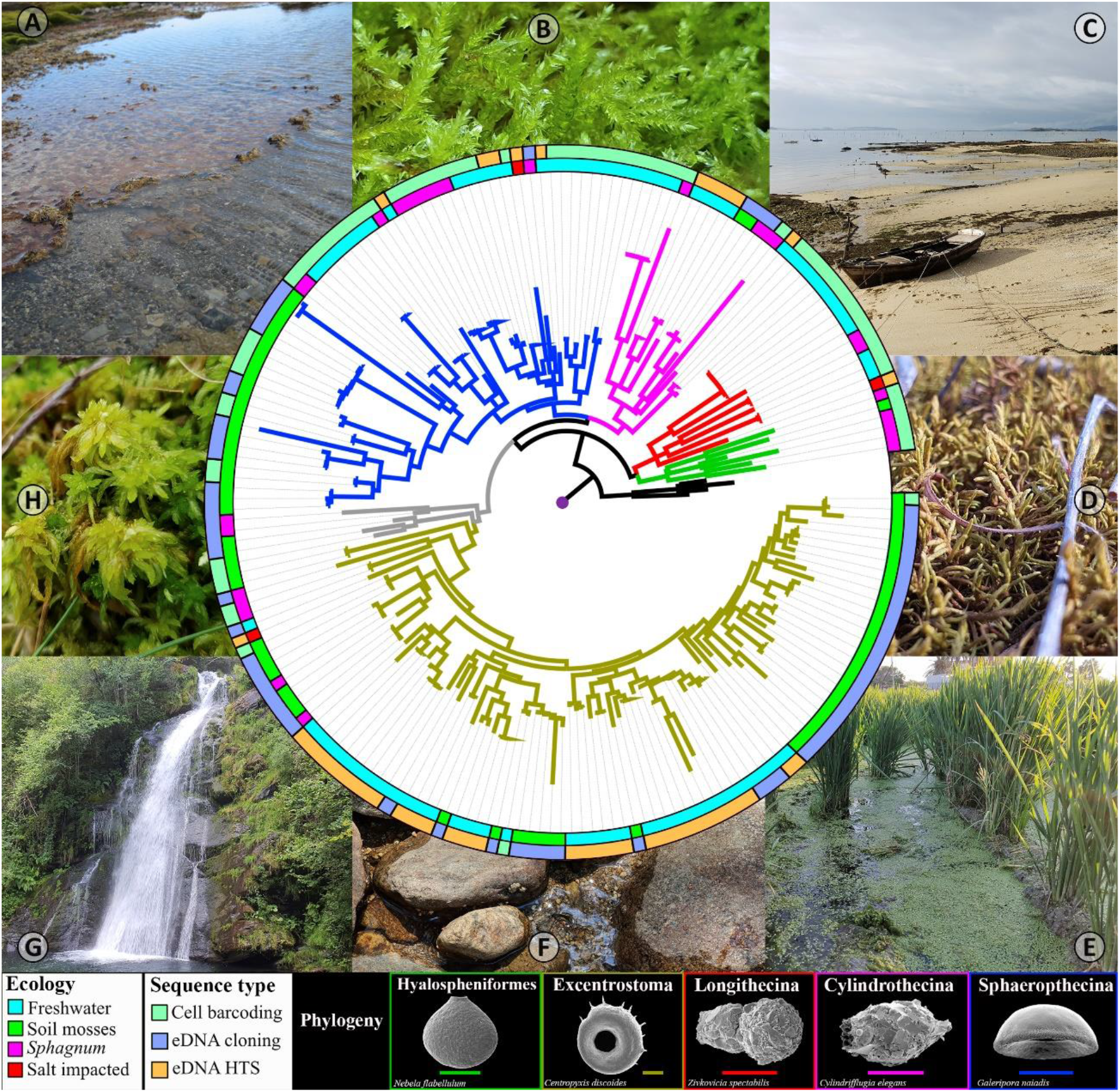
Maximum Likelihood phylogenetic tree of Arcellinida based on COI, with the branch colours representing the infraorders in Glutinoconcha. The inner circle represents the ecosystem type from where organisms derive and the outer circle the origin of sequences. Pictures represents the localities where the samples were taken A) Antarctic, B) wet mosses, C) mosses near the sea, D) dry mosses, E) eutrophicated freshwater, F) sediment from lake, G) sediment from a freshwater river and H) *Sphagnum*.

Recent studies involving the application of molecular protocols have provided new insights on both the diversity and taxonomy of Arcellinida. For example, the systematic barcoding of single species with the variable mitochondrial marker COI revealed the existence of crypto taxa not identifiable through morphological analysis alone (González-Miguéns, Soler-Zamora, Villar-Depablo, et al., 2022; Kosakyan et al., 2012; Singer et al., 2015). Although morphologically (almost) indistinguishable these so-called “pseudocryptic” species do differ from each other according their ecological preferences (Singer et al., 2018) and their global distributions (Singer et al., 2019). The combination of barcoding, distributional and ecological evidence indicates that these taxa should be considered as true biological species (Lara et al., 2020). Given the difficulties with identification of some taxa based solely on their morphology, a molecular approach is required for the collection of reliable distributional assessment. In addition to reducing costs and handling time, the application of a metabarcoding protocol for Arcellinida biodiversity assessment allows to overcome the previously described identification issues.

In this study, we introduce a species-level protocol to characterize Arcellinida diversity based on eDNA samples. We describe an eDNA extraction and amplification protocol based on an Arcellinida-specific primer, using the mitochondrial barcode gene COI. Finally, we describe a bioinformatic pipeline to analyse the resulting data. As a case study, we applied this protocol to a collection of samples from the Western Palearctic region covering sites with naturally high (peatlands) and low diversity/density of Arcellinida (estuaries). We subsequently generated an environmental sequences database based on both classical cloning/sequencing as well as Illumina sequencing. We also compare our metabarcoding data with morphological observations to explore the capacities of our polyphasic methodology to draw conclusions on the environmental diversity of Arcellinida.

## 2 Materials and Methods

### 2.1 Sampling

For this study we collected 42 samples from Europe, North America and the Antarctic Peninsula. We sampled freshwater sediments, terrestrial mosses, bogs (*Sphagnum*) and marine (interstitial) environments (Table S1). Soil and *Sphagnum* samples were collected following (González-Miguéns, Soler-Zamora, Villar-Depablo, et al., 2022) with the mosses and sediment being immersed in a falcon tube with distilled water. The moss was then squeezed to release the protist community, which was allowed to collect in the bottom of the tube to concentrate the testate amoebae. One ml of this concentrate was collected for subsequent analysis. Freshwater substrate samples were collected using a sterile plastic Pasteur pipette or a syringe (when scuba diving), collecting only the top-most few ml of sediment. Again, only a one ml aliquot of this sample was used for subsequent analysis. Samples were fixed and conserved within LifeGuard Soil Preservation Solution (Qiagen) (in a proportion 1:1, with respect to the sample) or preserved at −20°C until DNA extraction (Mazei et al., 2015). All samples were collected with different and sterile plastic Pasteur pipettes and Falcon tubes to avoid cross contaminations. Samples from Frog and Oromocto lakes (New Brunswick, Canada) and Nantes, Kérouarg, Kergironé and Beg ar Graz estuaries (France) were sequenced using Illumina Novaseq and MiSeq platforms, respectively. The rest of the samples were sequenced by Sanger dideoxy method, after cloned the PCR amplicons (Table S1).

### 2.2 eDNA extraction and amplification

The sampling and laboratory works were performed inspired by the review of (Goldberg et al., 2016). Total eDNA was extracted using the DNeasy PowerSoil Pro Kit (Qiagen) and FastDNA Spin Kit for Soil (MP Biomedicals), following the instructions provided by the manufacturer. In each round of eDNA extraction, a negative control was performed to track potential contamination. The extracted eDNA was conserved at −20 °C until further processing. Then, we amplified a portion of the cytochrome oxidase subunit I (COI), using a two-step nested polymerase chain reaction (PCR) protocol specifically designed for Arcellinida:

1. The first PCR had a final volume of 11 μl containing: 1 μl of DNA template, 1 μl of distilled water, 1 μl of bovine serum albumin (BSA), 6 μl MyTaq Red DNA polymerase Mix (BioLine) and 1 μl of each primer (10 μmol). We used the universal COI primers pair developed by (Folmer et al., 1994) LCO 1490 (5’ GGTCAACAAATCATAAAGATATTGG 3’), as forward, and HCO 2198 (5’ TAAACTTCAGGGTGACCAAAAAATCA 3’) as reverse. The PCR cycling profile was as follows: initial denaturation at 96 °C for 5 min, followed by 40 cycles at 94 °C for 15 s; 40 °C for 15 s and 72 °C for 90 s and a final extension step at 72 °C for 10 min. The resultant fragment was 692 bp long.
2. The second PCR had a final volume of 21 μl containing 1 μl of product amplification of the first PCR, 6 μl of distilled water, 12 μl MyTaq Red DNA polymerase Mix (BioLine) and 1 μl of each primer (10 μmol). We used LCO 1490 as the forward primer and the Arcellinida-specific primer developed in (González-Miguéns, Soler-Zamora, Villar-Depablo, et al., 2022) ArCOIR (5’ CCACYNGAATGWGCTARAATACC 3’), as reverse, with the following PCR cycling profile. There was an initial denaturation at 96 °C for 5 min, followed by 40 cycles at 94 °C for 15 s, 55 °C for 15 s and 72 °C for 90 s, and a final extension step at 72 °C for 10 min. The derived fragment was 407 bp long. To multiplex these samples, we developed custom dual indexes with spacers primers LCO and ArCOIR. A list of these primers is included in Table S2.

Each PCR was replicated three times, and then all replicates were pooled together to mitigate the effects of the PCR biases. These PCRs were performed in a “PCR workstations” equipped with a UV hood and HEPA filter, using filter tips and negative-positive controls, to avoid and check for potential contaminations. We used 3 μL of each PCR reaction product and analysed them by electrophoresis on a 1% agarose gel. The PCR products of the second PCR were then purified, by band excision. Finally, we quantified the eDNA using a Qubit 3 Fluorometer, with dsDNA High Sensitivity (HS) assay kits (ThermoFisher) and stored at −20 °C.

### 2.3 Sequencing

First, we evaluated the specificity of the ArCOIR primer in eDNA samples by cloning the PCR products, based on 32 samples (Fig. 1; Table S1). As the incorporation of the insert in the vector is a stochastic process depending on the inserts proportion and size, we consider that cloning/Sanger sequencing is a good approach to test the specify of the primers prior to the application of HTS technologies. We used the pGEM^®^ T-Easy Vector System kit (Promega) following the manufacturer’s recommendations. We selected 35 to 40 recombinant clones per library and amplified the insert with the plasmid primers T7 (5’ TAATACGACTCACTATAGGG 3’) and SP6 (5’ GATTTAGGTGACACTATAG 3’) universal primers, using the following volumes: the colony without performing a DNA extraction step, 6 μl of distilled water, 12 μl MyTaq Red DNA polymerase Mix (BioLine) and 1 μl of each primer (10 μmol). The resulting PCR of the colonies were sequenced with Sanger dideoxy-nucleotides technology, by the company Macrogen (Madrid, Spain).

To test the robustness and the validity of our methods we used two different sequencing platforms and companies to sequences the amplicons. Samples from the Canadian lakes (FRO2, ORO2, Q1, Q2 and Q3) were sequenced using a NovaSeq 6000 (Illumina) by the company Novogene UK (Cambridge, United Kingdom). The samples from the French estuaries (13, 35, 48, 78 and 79) we were sequenced using MiSeq (Illumina) by the company ID-Gene Ecodiagnostics (Geneva, Switzerland). In both cases the company performed a purification and library, by adapter ligation, and generated paired-end 250 bp reads.

### 2.4 Data curation

The resulting cloning PCR sequences were checked and trimmed from cloning vector sequence and primer using the software Geneious Prime ver. 2019.0.4. We only considered sequences which were obtained from at least three independent colonies. To corroborate the identity of the sequences as Arcellinida, we performed blastn analysis (Altschul et al., 1990) against the GenBank and our own database (González-Miguéns, Todorov, et al., 2022).

The Illumina reads were curated using the following pipeline: 1) primers trimming and demultiplexing was done with Cutadapt ver. 2.8 (Martin, 2011), following the pipeline “Cutadat_Piepline.bash” provided as supplementary materials; 2) reads were ordered in the same direction using MAFFT ver. 7.490 (Katoh & Standley, 2013); 3) the resulting reads per sample were analysed with the dada2 R package (Callahan et al., 2016), following the tutorial of https://benjjneb.github.io/dada2/tutorial_1_8.html, with minor modifications. First, we visualized the quality profiles of the forward and reverse reads, to filter and trim them, truncating the reads after positions 210 and 200 for the forward and reverse, respectively. We dereplicated the reverse and forward reads, and then merged the paired reads, generating amplicon sequence variants (ASV). Then, we removed chimeric sequences. We only considered the ASV that were composed of at least ten reads to reduce the consequences of methodological errors such as “tag jumping” (Schnell et al., 2015). 4) for the taxonomic assignation we used VSEARCH ver. 2.14.1 (Rognes et al., 2016) using a curated database from (González-Miguéns, Todorov, et al., 2022), including COI sequences from GenBank public database (Amoebozoa, Archaeplastida, Excavata, Opisthokonta and SAR). The ASV with an identity value higher than 0.82 with the Arcellinida database, were checked in GenBank to verify the taxonomic assignation.

### 2.5 Phylogenetic analysis

The sequences of environmental clones and ASV that were assigned to Arcellinida were aligned with the Arcellinida COI sequences data of González-Miguéns, Todorov, et al. (2022), using the MAFFT auto algorithm (Katoh et al., 2002) as implemented in Geneious ver. 2019.0.4. Tree topologies and node supports were evaluated with maximum likelihood (ML), using IQ-TREE2 ver. 2.0 (Minh et al., 2020). The best substitution models were selected with ModelFinder (Kalyaanamoorthy et al., 2017). Node supports were assessed with 5000 ultrafast bootstrap replicates approximation (Hoang et al., 2018; Minh et al., 2013). The trees obtained were edited in FigTree v.1.4.3. Scanning electron microscopy images in the figures were taken from Todorov & Bankov (2019).

To visualize the sequence patterns in the region of the ArCOIR primer we used the ggseqlogo R package (Wagih, 2017) to create sequence logos for each Glutinoconcha infraorder. We retrieved the sequences from single cell from (González-Miguéns, Todorov, et al., 2022) and the Hyalospheniformes GenBank database, extracting the ArCOIR primer region.

### 2.6 Morphology vs molecular

We compared our molecular data with Arcellinida biodiversity assessments based on morphology from the same Canadian lake’s samples (Steele et al., 2018). This study was based on a total of 43 samples, divided in three quadrants “Q1”, “Q2” and “Q3”. Our samples “Q1”, “Q2” and “Q3”, sequenced by Illumina, correspond with one sample per quadrant in Steele et al. (2018), labelled with the same name.

To compare Arcellinida morphotypes retrieved in Steele et al. (2018) with our molecular ASVs, we clustered them into Operational Taxonomic Units (OTU). We consider as an OTU grouping the sequences based on the uncorrected pairwise distances (“p”) of 5% of genetic divergence. The OTUs composition of the samples Q1-3 was compared with the morphospecies recovered in Steele et al. (2018) for each family recovered in the study (Table S3). As the families Difflugidae and Cylindrifflugidae are practically impossible to differentiate morphologically from each other (González-Miguéns, Todorov, et al., 2022) we merged these taxa within a family “Difflugiidae”. In order to evaluate the “coverage” of Morphology vs Molecular we evaluate the coverage (C) as the ratio between the number of morphospecies retrieved in Steele et al. (2018) and the number of molecular OTUs (morphological OTUs/molecular OTUs). Where C=1 if the number of morphospecies is equal to the number of molecular OTUs, C>1 if morphological diversity is higher than the assayed molecular one and C<1 if molecular diversity is higher than morphological.

## 3 Results

### 3.1 Molecular results

We obtained a total of 82 Arcellinida clonal sequences, each of them being recovered in at least three independent colonies per sample (Fig. 1 and Table S1). In all environments surveyed (freshwater sediments, soil and bogs), Arcellinida clonal sequences dominated together with a minor representation of sequences classified as diatoms or naked amoebae.

The number of Illumina reads per sample is detailed in (Table S4), as well as the number of reads assigned to Arcellinida (Fig. 2 and Table S5). We obtained in total 61 ASV classified as Arcellinida. The samples from freshwater lake sediment provided the highest number of Arcellinida 73%, 91%, 48%, 67% and 76% from the total of reads, in the samples FRO2, ORO2, Q1, Q2 and Q3, respectively. The samples from estuary sediment provided low numbers of Arcellinida reads: 1%, 0.08%, 0.03%, 0.04% and 0.1% from the total of reads, in the samples 79, 78, 48, 35 and 13, respectively. The best represented eukaryotic clade outside Arcellinida was Lobosa “naked amoebae” (Fig. 2), with 25%, 7%, 41%, 23% and 21% from the total of reads, in the samples FRO2, ORO2, Q1, Q2 and Q3, respectively (freshwater sediments), This was also the case in estuary sediment samples with 95%, 7%, 96%, 26% and 78% from the total of reads, in the samples 79, 78, 48, 35 and 13, respectively.

**Figure 2.**
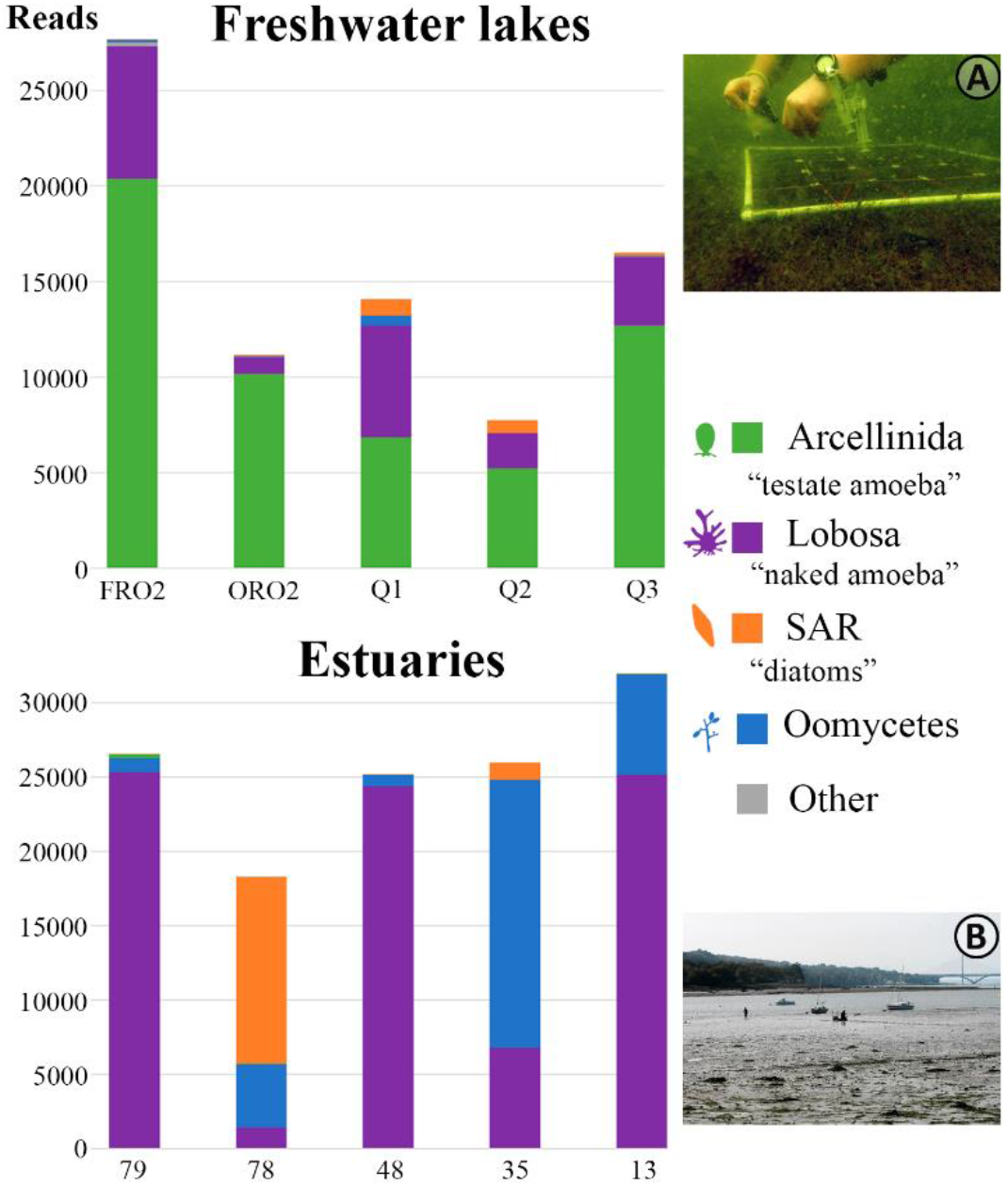
Number of reads recovered for each sample sequenced by Illumina. The colour represents the taxonomic composition. The pictures illustrate the type of environment in which samples were taken A) Q1, Oromocto Lake (Canada) and B) 13, Elorn (France).

### 3.2 phylogenetic composition of communities

In total we obtained 143 environmental sequences assigned to Arcellinida by combining cloning/sequencing and Illumina (Table S1). These sequences were placed in a phylogenetic tree whose topology was congruent with previous works (González-Miguéns, Todorov, et al., 2022) (Fig. 1). Notably, we recovered the monophyly of Arcellinida with a maximum likelihood bootstrap (ML) of 100 (Fig. 3). The tree also recovered the monophyly of the Glutinoconcha infraorders Hyalospheniformes ML= 100 (green colour in Fig. 1 and 3), Longithecina ML= 98 (red colour in Fig. 1 and 3), Sphaerothecina ML= 74 (blue colour Fig. 1 and 3), Cylindrothecina ML= 91 (purple colour in Fig. 1 and 3) and Excentrostoma ML= 92 (yellow colour in Fig. 1 and 4). Two clades did not classify among these infraorders, one of them also including the characteristic species *Trigonopyxis arcula;* we labelled them Arcellinida *incertae sedis* (grey colour in Fig. 1 and 4).

**Figure 3.**
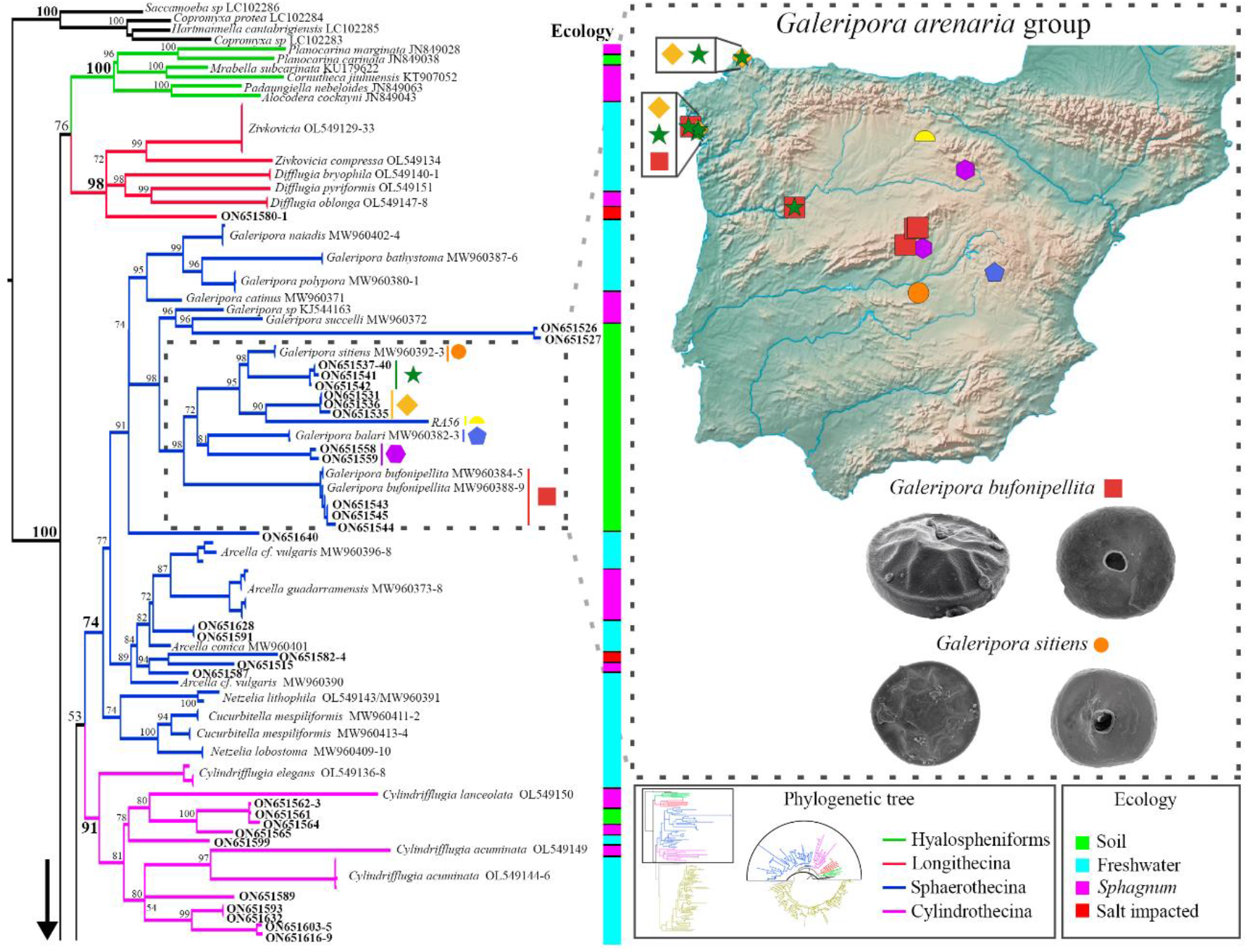
Detail of the phylogenetic tree of Arcellinida based on COI (Fig. 1). The symbols on the map of the Iberian Peninsula represent the different localities where sequences of the group of *Galeripora arenaria* species complex have been found. At the bottom are SEM images for *Galeripora bufonipellita* and *G. sitiens*, with typical habitat for each of the species.

**Figure 4.**
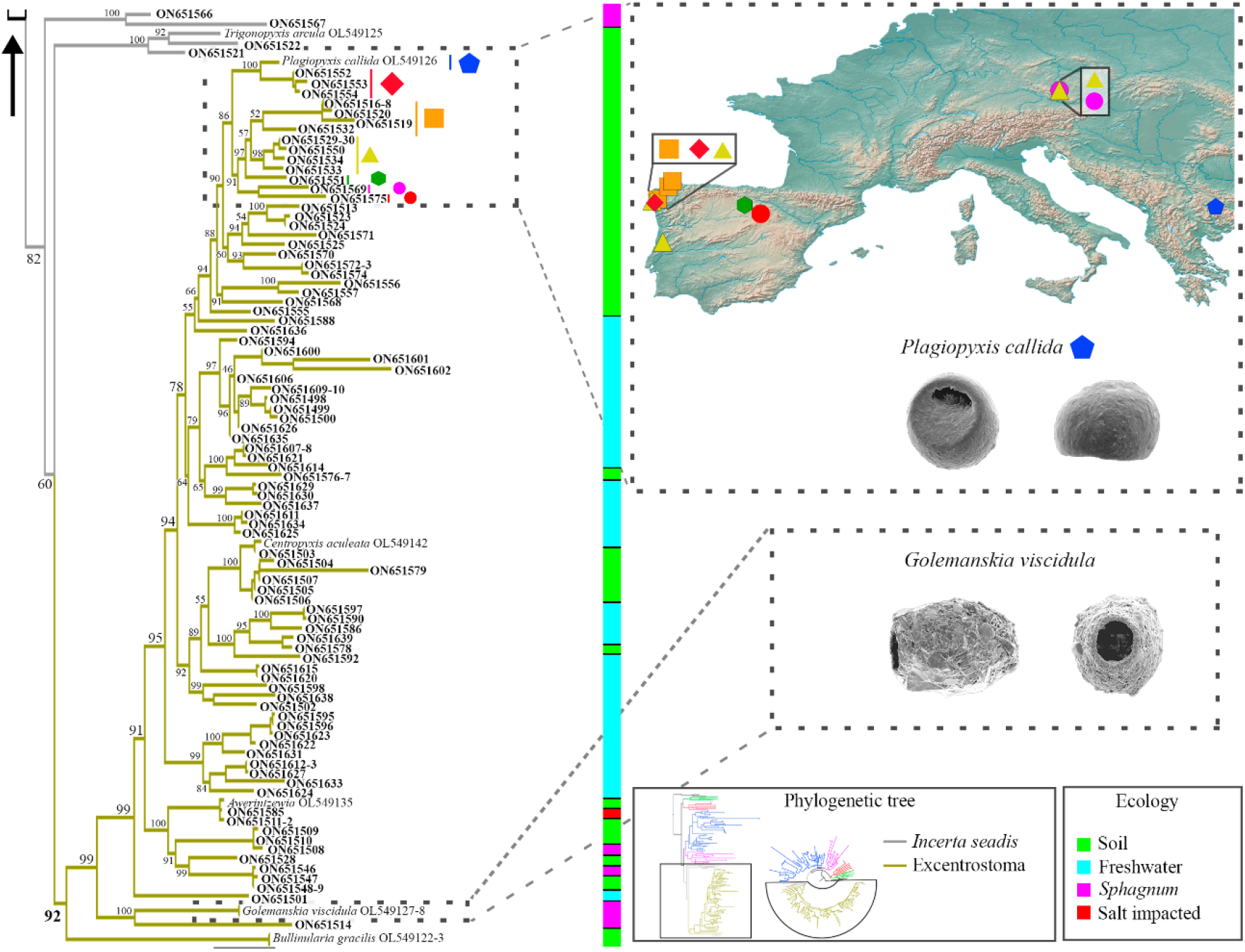
Detail of the phylogenetic tree of Arcellinida based on COI (Fig. 1). Symbols on the map of Europe represent the different localities where species from the *Plagiopyxis*clade have been found. At the bottom are SEM images for *Plagiopyxis callida* and *Golemanskia viscidula*, with typical habitat for the species.

Environmental sequences were distributed among Glutinoconcha infraorders in the following way: Hyalospheniformes 0 (0 cloned (C) and 0 Illumina (I)), Longithecina 1 (0 C and 1 I), Sphaerothecina 20 (15 C and 5 I), Cylindrothecina 10 (4 C and 6 I), Excentrostoma 86 (48 C and 38 I) and *Incertae Sedis* 4 (4 C and 0 I) (Fig 1,3 and 4). Some of these ASV corresponded to organisms that had been previously isolated and barcoded (González-Miguéns et al., 2022a; 2022b). These taxa were *Galeripora bufonipellita, Centropyxis aculeata* and *Awerintzewia* sp.

Sequences from Hyalospheniformes were not recovered, even though these organisms could be observed microscopically in the samples. This was probably because these organisms have a codon deletion at the beginning of the primer (Fig. 5). The low coverage of Longithecina is also probably due to a mismatch of their COI sequence with the primer ArCOIR (Fig. 5). Reference sequences of the infraorders Volnustoma (within Glutinoconcha) and of the suborders Organoconcha and Phryganellina are not yet barcoded. The expansion of the barcode database will allow finding the limits of our protocol at deeper taxonomic levels.

**Figure 5.**
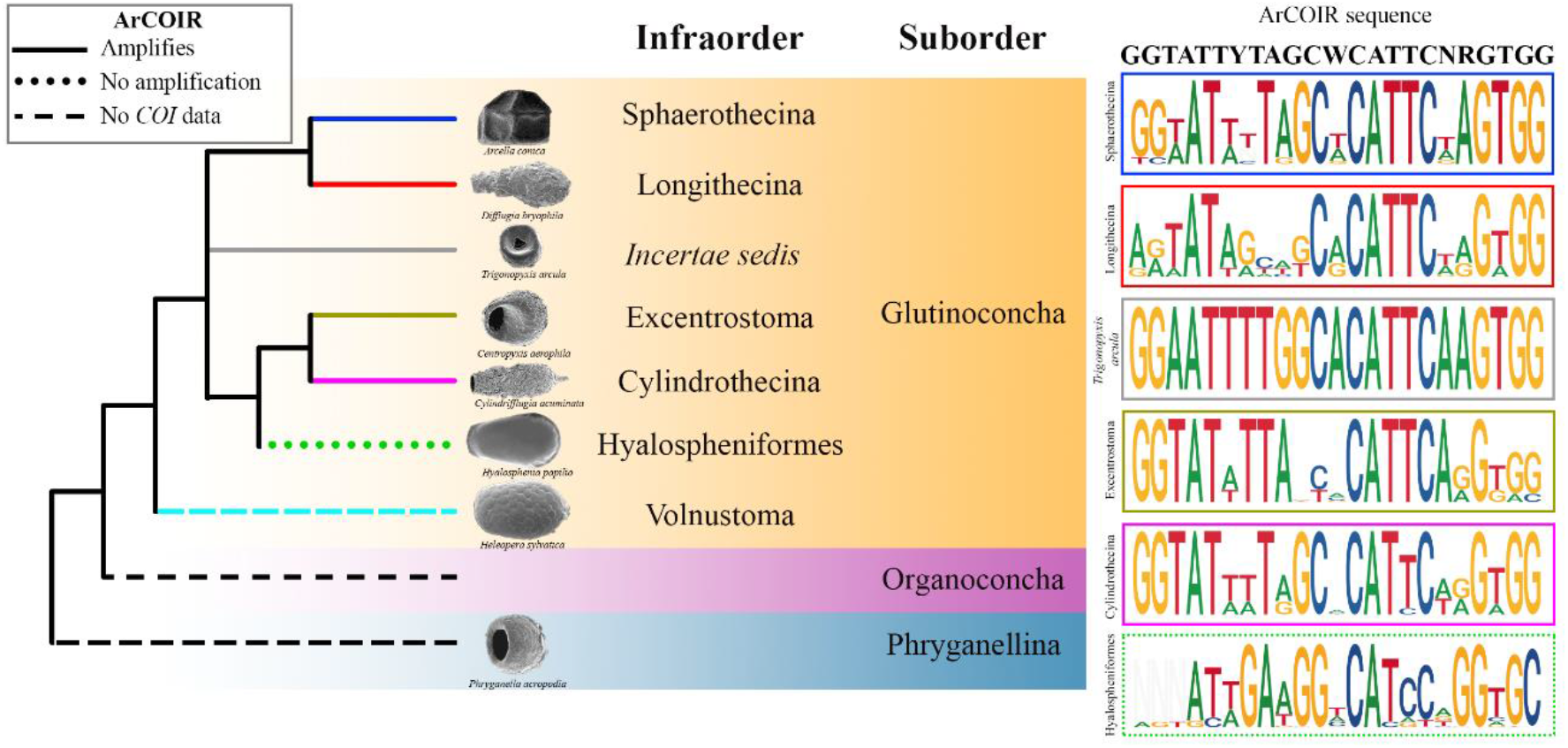
Schematic phylogenetic tree of Arcellinida based on the results of (González-Miguéns, Todorov, et al., 2022; Lahr et al., 2019). The branch style illustrates if members of the infraorder can be amplified with our newly designed primer ArCOIR or not. On the right panel, sequence logos represents the base ratio, in the ArCOIR primer region, based on single cell sequences.

### 3.3 Morphology vs molecular results

Calculation of coverage can provide hints regarding the potential and any limitations of our metabarcoding protocol by comparing both morphological classification in addition to the molecular data, for each clade. Even though the sampling intended for metabarcoding was much smaller than the one in Steele et al., 2018, we observed that some clades were better represented in metabarcoding than by direct counts. Indeed, the families with less coverage (better represented in molecular data than in microscopic counts), were Arcellidae C= 0 (no morphological observations) and Centropyxididae C= 0.57. On the other hand, the families with better morphological coverage were Difflugiidae C= 5 and Netzeliidae C= - (not recovered in the molecular data). The C values for all taxa with respect to (Steele et al., 2018) are given in Table 1.

**Table 1.**
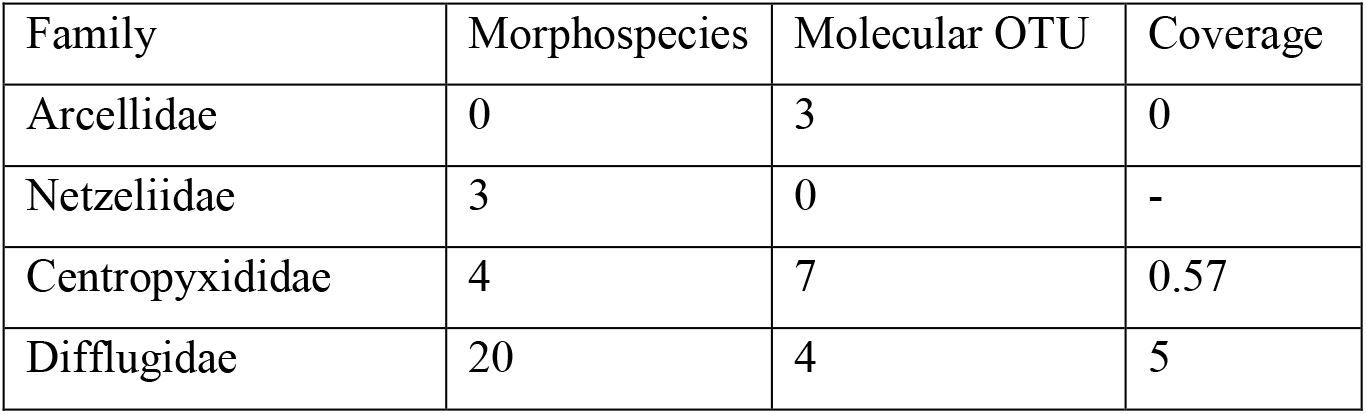
Coverage between Morphospecies and Molecular OTU

## 4 Discussion

### 4.1 Prerequisites for species level protocols in eDNA

As species are the only biologically and ecologically meaningful (and comparable) units of diversity, species-level resolution in metabarcoding studies has many advantages. Cross-taxa comparisons can be generated including information about each single species such as distribution, ecological optimum and tolerance or ecological function. One of the requisites for a “resolutive” marker gene must be variable enough to reflect the evolutionary history of each taxonomic unit, as each species has its own historical and future evolutionary trajectory with respect to other clades (Wiley, 1978). In Arcellinida, these barcodes were integrated with shell morphology (Singer et al., 2015) ecology (Singer et al., 2018), biogeography (Singer et al., 2019), or with several molecular markers (González-Miguéns, Todorov, et al., 2022), to trace species delimitation. The mitochondrial cytochrome oxidase subunit I (COI) was shown to be the best adapted marker for species-level discrimination, in Arcellinida but also in other Amoebozoa (Nassonova et al., 2010; Tekle, 2014). This gene is, to date, one of the most used barcoding genes in general (except for plants) for species identification, and also for species level metabarcoding (Bucklin et al., 2021; Creedy et al., 2021).

The fine taxonomic resolution reached with our eDNA protocol, based in COI, makes clearer the evaluation of biodiversity patterns such as ecosystem evolution (Pawlowski et al., 2018) which may act on functional diversity as well (Laroche et al., 2021). Because eDNA facilitates the collection of high amounts of species molecular data with minimal effort, it represents an ideal approach to these applications, as well as to increase the existing knowledge of the ecology or distribution of species. These practices are much more common in plants and animals, and bring along many applications, for instance to invasive species monitoring (Ficetola et al., 2008), food regime studies (Roffler et al., 2021) and even to the rediscovery of critically endangered species (Turunen et al., 2021; Villacorta-Rath et al., 2022). The diversity of these applications suggests the immense potential that lies in the application of species level metabarcoding to new protist groups.

Regarding to microorganisms, eDNA is commonly the only way to retrieve information from elusive species, without direct observation (Pawlowski et al., 2012). Often, however, sequences retrieved are not sufficient to draw any biologically relevant conclusions, as no close-related species have been entered in the database. These sequences resulting from lack of taxonomic information on microbial eukaryotic diversity form so-called “environmental clades”; whose number and importance depend on the degree of species single cells barcoding effort. In our data, we observed only one single group, which cannot be assigned yet to any known Glutinoconcha infraorder. We designated this group as *incertae sedis*, pending morphological characterization of these organisms (Fig 2). Here, we provide the most complete COI database, which includes all infraorders described as for now within Glutinoconcha, except Volnustoma and Hyalospheniformes, which are soil-dwelling lineages. While we do not possess sequences from Volnustoma, many Hyalospheniformes typically have a codon deletion on the binding site of the primer ArcCOIr (Fig. 5). This concerns the clade which includes the sister clade of the rest of Hyalospheniformes and comprises genera *Alocodera, Apodera* and *Padaungiella*.

### 4.2 Implications on the current knowledge of Arcellinida’s environmental diversity

The application of our metabarcoding protocol has challenged previous perceptions of Arcellinida diversity. The most conspicuous outcome is the immense diversity of infraorder Excentrostoma, which includes the COI-barcoded genera *Centropyxis, Plagiopyxis, Bullinularia, Awerintzewia* and the recently described genus *Golemanskia* (Fig. 1 and 4). This order includes a vast diversity of shell morphologies (González-Miguéns, Todorov, et al., 2022) well-beyond the typical compressed, off-centered apertured shell (Lahr et al., 2019). Here, we retrieved an immense diversity in all environments surveyed (including in estuaries), suggesting an unexpected diversity overall (Fig. 1 and 4). As might be expected the diversity of Arcellinida observed was closely related to the environment they were found in. For instance, *Galeripora arenaria* complex species are only recorded in soil (Fig. 3) as observed with the *Plagiopyxis* group (Fig. 4). On the other hand, several other well supported environmental groups belonged exclusively in aquatic sediments (Fig. 4). This result suggests a widespread occurrence of phylogenetic niche conservatism that should be tested. This trend has been demonstrated previously in Hyalospheniformes, where tolerance towards nitrogen depletion in *Sphagnum* peatbogs is inherited within clades in the genus *Nebela* (Singer et al., 2018).

It appears obvious, comparing the presence of Arcellinida in freshwater versus estuary sediments, that the first contain the highest diversity and most probably the highest individual numbers (Fig. 2). Estuaries, because of their fast changing salinity and their dynamic nature, contain generally a low diversity for most taxa, including the macroscopic ones (Benfield, 2012). Arcellinida have been recorded only on a limited number of occasions in high salinity environments (Smith et al., 2009); they typically become scarcer in anchialine systems as the salinity increases (Van Hengstum et al., 2008). It seems therefore that Arcellinida species are primarily a freshwater taxon. However, we recovered an ASV in the estuarine sediment which corresponds exactly to *Awerintzewia* sp., an isolate obtained from salt impacted bryophytes growing on a beach. The same sequence was also found again in a clone from a similar environment (Fig. 4). This suggests a trend towards halophily for this species, and that some Arcellinida at least are capable of crossing the salinity barrier and establish populations in higher salinity environments. Two other ASV were also recovered in estuarine sediment, corresponding respectively to a member of genus *Arcella* as well as an unidentified Longithecina. The salinity barrier is one of the greatest ecological barriers, and those organisms that cross it must face important challenges through their evolutionary history. This characteristic has been shown in another group of testate amoebae, *Cyphoderia* (González-Miguéns, Soler-Zamora, Useros, et al., 2022). Therefore, it can be expected that further investigations in saline environments, both inland (saline endorheic) and marine systems will reveal new insights on Arcellinida diversity.

### 4.3 morphological and molecular integration in Arcellinida’s applied studies

The comparison between morphological observations and OTUs (see morphology Vs molecular in Materials and Methods) through coverage indices reveals the great potential of our new approach for diversity exploration (Table 1). The families Arcellidae and Centropyxididae had low coverage indices (0 and 0.57, respectively, in Table 1), meaning that metabarcoding retrieved more diversity than the morphological observations in Steele et al., 2018. This was even more remarkable as our sampling design is much more reduced (3 versus 43 samples) and the size of our samples was smaller (ca. 1 cm^3^ of unprocessed sediment versus about 5 cm^3^ of concentrated sediment). Several explanations can be invoked to explain these differences between visual observations and metabarcoding results. Arcellinida have inconspicuous “naked states” in their lifecycles which are very difficult to observe within environmental samples (Dumack et al., 2020). In addition, many Arcellinida species are difficult to distinguish visually but are genetically different (cryptic diversity). The difficulty in species identification in Arcellinida, has already been observed by Foissner & Korganova (2000) when they hypothesized that *Centropyxis aerophila* was a cryptic species complex. These authors, by observing the immense morphological diversity continuum of the group, already hypothesized a diversity higher than expected. For practical reasons, these forms were pooled together in ecological studies into a “*Centropyxis aerophila*-type (Amesbury et al., 2018).

In contrast, the family Difflugiidae shows a low coverage index (5 in Table 1), and no sequence has been detected for the family Netzeliidae. Sample size certainly plays a role, as *Cucurbitella mespiliformis* has been detected visually but not by metabarcoding. Indeed, the primer ArcCOIr recognized only few mismatches with that species (Fig. 5). Primer incompatibility may play a role in “Difflugiidae”, by reducing the likelihood of their COI genes being amplified during the PCR process (Fig. 5). Indeed, several mismatches with ArCOIR can be found in some members of genus *Difflugia*, while this is less of an issue in *Cylindrifflugia* (Fig 1 and 2). Then, Arcellinida populations density are known to vary during the year, as their metabolic activity and reproduction rates are influenced by water temperature as shown for *Netzelia tuberspinifera* in the transcriptome-based study of Wang et al. (2022). A larger sampling design with repeated sessions along a full seasonal cycle should allow retrieving environmental sequences from the less represented families in this study.

## 5. Conclusion

A well-designed metabarcoding protocol will recover most of the target diversity present in an environment. Our new Arcellinida metabarcoding protocol recovers all the infraorders in Glutinoconcha, with COI data (except for Hyalospheniformes) at the species level. It will allow for the rapid, cost effective and reliable handling of data in potentially vast studies for the monitoring of ecosystem health, as well as ecological or biogeographic data from samples collected all over the world. The application of Arcellinida, an important mid trophic bioindicator can fill the current monitoring gap in lentic (e.g. lakes and reservoirs) and soil ecosystems. Further studies that include COI barcoding of additional documented organisms will improve the interpretative power of Arcellinida metabarcoding studies, and for example provide a deeper understanding of many areas of research with the group, such as the link between Arcellinida test traits and ecological functions (Marcisz et al., 2020). Altogether, this information will contribute to make Arcellinida an ideal model group for theoretical and applied studies.

## Acknowledgements

We express our gratitude to C. Soler-Zamora, F. Useros, Dr I. García-Cunchillos and M. Blázquez for helpful during the manuscript. We also acknowledge the help of Y. Ruiz-León and M. García-Gallo for the help in the molecular biology laboratory and M. Villar-De Pablo for the data processing and analyses. We are indebted to the following people that took part in the field work; A. Berlinches, A. Kosakyan, I. Cacabelos, Prof. E. A. D. Mitchell, L. Hernandez, L. Lara-Hollenstein, M. Blázquez, M. Miguéns-Gomez, M. Villar-De Pablo, P. Lara-Hollenstein. We acknowledge Prof F. Jorissen and the project FORESTAT (funded by the OFB (Office français de la biodiversité) and the University of Angers, grant number 3976-CT_RD_AMI_18_SURV_FORESTAT) for the estuarine samples. This study was funded by the Spanish Government PGC2018-094660B-I00 /10.13039/501100011033/ (MCIU/AEI/FEDER,UE) and an “Atracción de Talento Investigador” grant awarded by the Consejería de Educación, Juventud y Deporte, Comunidad de Madrid (Spain) (2017-T1/AMB-5210) to E.L.; a National Geographic Grant EC-86022R-21 to R. G. M. Field work in New Brunswick Canada was funded by a Natural Sciences and Engineering Research Council (NSERC) of Canada Discovery Grant to R.T.P.

