## Supplementary material for "A needle in a haystack: a new metabarcoding approach to survey diversity at the species level of Arcellinida (Amoebozoa: Tubulinea)": Table s1

**Short title**: metabarcoding species level for Arcellinida

Rubén González-Miguéns^1,*^, Emilio Cano^1, 2^, Antonio Guillén-Oterino^3^, Antonio Quesada^4^, Daniel J.G. Lahr^5^, Daniel Tenorio-Rodríguez^3^, David de Salvador-Velasco^3^, David Velázquez^4^, María Isabel Carrasco-Braganza^3^, R. Timothy Patterson^6^, Enrique Lara^1,*^, David Singer^6,7^

**Table of Contents:**

| **Table S1** | Pages 1-6 |
| --- | --- |
| **Table S2** | Page 7 |
| **Table S3** | Page 8 |
| **Table S4** | Page 9 |
| **Table S5** | Page 10 |

**Table S1**. Localities sampled, with information of the habitat and GenBank accession of the sequences retrieved in each locality.

| **ID** | **locality** | **coordinates** | **ecology** | **date** | **GenBank** |
| --- | --- | --- | --- | --- | --- |
| **Cloned sequences** | | | | | |
| R2 | **Antarctic**: Byers Peninsula | -62.6272, -61.0230 | **Freshwater**: near river | 2019 | ON651501 |
| R23 | **Czech Republic**: Ceske Budejovice, park Stromovka | 48.969444, 14.458611 | **Soil**: mosses | 2019 | ON651533, ON651548, ON651568, ON651569 |
| R24 | **Czech Republic**: Ceske Budejovice, park Stromovka | 48.969444, 14.458612 | **Soil**: mosses, near pond walls | 2019 | ON651562, ON651549 |
| R29 | **France**: Frasne, La Tourbière | 46.832067, 6.157517 | ***Sphagnum***: wet part | 2019 | ON651566, ON651567 |
| R6 | **France**: Frasne, La Tourbière | 46.832067, 6.157517 | ***Sphagnum***: dry part | 2019 | ON651508, ON651546, ON651565, ON651563 |
| R17 | **Portugal**: Coimbra, rúa Inácio Duarte | 40.210556, -8.424678 | **Soil**: mosses in a tree | 30/06/2019 | ON651529, ON651534, ON651557 |
| R18 | **Portugal**: Coimbra, jardim Sereia | 40.209090, -8.417197 | **Soil**: mosses in a tree | 30/06/2019 | ON651530 |
| R1 | **Spain**: Zamora, Lake Sanabria | 42.120420, -6.731126 | **Freshwater**: 3 m deep | 2019 | ON651498, ON651499, ON651500, ON651502 |
| R3 | **Spain**: A Coruña, playa de Cariño | 43.732840, -7.871843 | **Soil**: mosses near the sea | 14/08/2020 | ON651503, ON651531, ON651532, ON651537, ON651536 |
| R4 | **Spain**: Burgos | 42.344733, -3.706544 | **Soil**: mosses | 16/02/2020 | ON651504, ON651507, ON651509, ON651556, ON651571, ON651579, ON651570 |
| R5 | **Spain**: Burgos | 42.343656, -3.703844 | **Soil**: wall mosses | 16/02/2020 | ON651505, ON651560, ON651551 |
| R7 | **Spain**: Burgos, Covanera | 42.735664, -3.801736 | **Soil**: mosses near river | 15/02/2020 | ON651510 |
| R8 | **Spain**: A Coruña, Cariño, mirador de Miranda | 43.716821, -7.896318 | **Soil**: mosses | 14/08/2020 | ON651511, ON651526, ON651527, ON651528 |
| R9 | **Spain**: Pontevedra, O Grove | 42.478333, -8.901417 | **Soil**: mosses near the sea | 2020 | ON651538, ON651512, ON651541, ON651552, ON651553, ON651554 |
| R10 | **Spain** |  | **Soil**: mosses |  | ON651513 |
| R11 | **Spain**: Zamora, Laguna de los Peces | 42.172030, -6.729745 | ***Sphagnum*** | 2019 | ON651514 |
| R12 | **Spain**: A Coruña, Fragas Eume | 43.423889, -8.125556 | ***Sphagnum*** | 15/08/2020 | ON651515 |
| R13 | **Spain**: A Coruña, Fragas Eume | 43.426011, -8.128678 | **Soil**: mosses | 15/08/2020 | ON651516, ON651520 |
| R14 | **Spain**: Pontevedra, Catoira | 42.687019, -8.726286 | **Soil**: mosses | 7/03/2020 | ON651517, ON651524, ON651525 |
| R15 | **Spain**: Pontevedra, Catoira | 42.668294, -8.714286 | **Soil**: mosses on a wall | 12/07/2019 | ON651518, ON651519 |
| R16 | **Spain**: Asturias, Cudillero, playa del Silencio | 43.569389, -6.293333 | **Soil**: mosses near the sea | 14/08/2020 | ON651521, ON651522, ON651523 |
| R19 | **Spain**: Pontevedra, Illa de Arousa, Carreiron | 42.531011, -8.869114 | **Soil**: mosses near the sea | 8/03/2020 | ON651539, ON651535 |
| R20 | **Spain**: A Coruña, Corrubedo, faro | 42.5755, -9.0898 | **Soil**: mosses | 13/08/2020 | ON651540 |
| R21 | **Spain**: Salamanca, Aldeadávila de la Ribera | 41.211111, -6.677039 | **Soil**: wall mosses, near a fontain | 22/02/2020 | ON651576, ON651542 |
| R22 | **Spain**: Soria, Playa Pita | 41.852589, -2.783518 | **Soil**: leaf litter near a lake | 11/05/2019 | ON651577, ON651575, ON651558 |
| R25 | **Spain**: A Coruña, Corrubedo | 42.584258, -9.045280 | **Soil**: dry mosses | 13/08/2020 | ON651543, ON651544, ON651547, ON651550 |
| R26 | **Spain**: Salamanca, Aldeadávila de la Ribera | 41.213872, -6.676036 | **Soil**: wet mosses | 22/02/2020 | ON651545 |
| R27 | **Spain**: Pontevedra, O Grove | 42.470972, -8.881361 | **Soil**: mosses | 2020 | ON651555 |
| R28 | **Spain**: Madrid, Fuencarral-El Pardo | 40.509789, -3.751213 | **Soil**: mosses in a tree | 20/07/2019 | ON651559, ON651561, ON651564 |
| R30 | **Spain**: Galicia, Pontevedra, Catoira | 42.686717, -8.727122 | **Soil**: mosses in a tree | 7/03/2020 | ON651572 |
| R31 | **Spain**: Lugo, Guitiriz, río Vega | 43.186386, -7.890778 | **Soil**: wet mosses | 29/02/2020 | ON651573, ON651574 |
| R32 | **Spain**: Ávila, Guisando | 40.223453, -5.151967 | **Soil**: wet mosses, near a river | 4/08/2020 | ON651578 |
| **ASV from Illumina** | | | | | |
| **Canadian lakes** | | | | | |
| FRO2 | **Canada**: Frog lake | 45.649632, -67.036067 | **Freshwater**: lake sediment | 19/08/2019 | ON651586, ON651590, ON651594, ON651597, ON651598, ON651600, ON651601, ON651602, ON651606,  ON651609, ON651611, ON651620, ON651622, ON651623, ON651624, ON651625, ON651629, ON651631, ON651634, ON651635, ON651639 |
| ORO2 | **Canada**: Wightman Cove Oromocto Lake | 45.6428, -66.9949 | **Freshwater**: lake sediment | 19/08/2019 | ON651607, ON651615, ON651616, ON651621, ON651630, ON651636, ON651637, ON651638 |
| Q1 | **Canada**: Wightman Cove Oromocto Lake | 45.641127, -66.999987 | **Freshwater**: lake sediment, water depth (3 m) | 18/08/2019 | ON651591, ON651593, ON651603, ON651612, ON651617, ON651640 |
| Q2 | **Canada**: Wightman Cove Oromocto Lake | 45.641914, -66.997589 | **Freshwater**: lake sediment, water depth (5 m) | 18/08/2019 | ON651588, ON651592, ON651595, ON651596, ON651604, ON651618, ON651632, ON651633 |
| Q3 | **Canada**: Wightman Cove Oromocto Lake | 45.642866, -66.994968 | **Freshwater**: lake sediment, water depth (6 m) | 18/08/2019 | ON651587, ON651589, ON651599, ON651605, ON651608, ON651610, ON651613, ON651614, ON651619, ON651626, ON651627, ON651628 |
| **French estuaries** | | | | | |
| 79 | **France**: Nantes | 47.2173, -1.5098 | **Marine**: subtidal sediment | 5/02/2021 | ON651582 |
| 78 | **France**: Nantes | 47.2069, -1.7305 | **Marine**: Subtidal sediment | 5/02/2021 | ON651583 |
| 48 | **France**: Auray, Kerouarch | 47.584956, -2.9608467 | **Marine**: Intertidal mudflat | 16/09/2020 | ON651580, ON651584 |
| 35 | **France**: Crac’h, Kerguironé | 47.610081, -3.024608 | **Marine**: Intertidal mudflat | 20/10/2020 | ON651581 |
| 13 | **France**: Elorn, Beg ar Graz | 48.424608, -4.304950 | **Marine**: Intertidal mudflat | 17/10/2020 | ON651585 |

**Table S2**. List of forward and reverse primers with barcode for multiplexing, in red the barcode and in purple the spacer.

| **LCO** | **ArCOIR** |
| --- | --- |
| NNACACACACGGTCAACAAATCATAAAGATATTGG | NNNNCAGAGACGCCACYNGAATGWGCTARAATACC |
| NNNACGACTCTGGTCAACAAATCATAAAGATATTGG | NNCAGATGATCCACYNGAATGWGCTARAATACC |
| NNNNACGCTAGTGGTCAACAAATCATAAAGATATTGG | NNNCAGTATGACCACYNGAATGWGCTARAATACC |
| NNACTATCATGGTCAACAAATCATAAAGATATTGG | NNNNCATAGTATCCACYNGAATGWGCTARAATACC |
| NNNACTGCTGAGGTCAACAAATCATAAAGATATTGG | NNCATGTGCTCCACYNGAATGWGCTARAATACC |
| NNNNAGACATCTGGTCAACAAATCATAAAGATATTGG | NNNCGAGAGATCCACYNGAATGWGCTARAATACC |
| NNAGTCTACAGGTCAACAAATCATAAAGATATTGG | NNNNCGAGTACGCCACYNGAATGWGCTARAATACC |
| NNNCAGATCACGGTCAACAAATCATAAAGATATTGG | NNCGATGTAGCCACYNGAATGWGCTARAATACC |
| NNNNCATACTGCGGTCAACAAATCATAAAGATATTGG | NNNCGTATAGACCACYNGAATGWGCTARAATACC |
| NNCATATACTGGTCAACAAATCATAAAGATATTGG | NNNNCGTGATGTCCACYNGAATGWGCTARAATACC |
| NNNCATCATATGGTCAACAAATCATAAAGATATTGG | NNGAGATAGTCCACYNGAATGWGCTARAATACC |
| NNNNCGACTCATGGTCAACAAATCATAAAGATATTGG | NNNGAGTGTCTCCACYNGAATGWGCTARAATACC |
| NNCGAGACGCGGTCAACAAATCATAAAGATATTGG | NNNNGCATATATCCACYNGAATGWGCTARAATACC |
| NNNCGAGCACAGGTCAACAAATCATAAAGATATTGG | NNGCATGACGCCACYNGAATGWGCTARAATACC |
| NNNNCGTATCGAGGTCAACAAATCATAAAGATATTGG | NNNGCGAGTAGCCACYNGAATGWGCTARAATACC |
| NNTAGACAGTGGTCAACAAATCATAAAGATATTGG | NNNNGCTATGATCCACYNGAATGWGCTARAATACC |
| NNNTAGCACGAGGTCAACAAATCATAAAGATATTGG | NNGCTGATCGCCACYNGAATGWGCTARAATACC |
| NNNNTATGACACGGTCAACAAATCATAAAGATATTGG | NNNTAGTAGAGCCACYNGAATGWGCTARAATACC |
| NNTATGTAGAGGTCAACAAATCATAAAGATATTGG | NNNNTATGATAGCCACYNGAATGWGCTARAATACC |
| NNNTCAGCTACGGTCAACAAATCATAAAGATATTGG | NNTCAGTAGTCCACYNGAATGWGCTARAATACC |
| NNNNTCAGTCGTGGTCAACAAATCATAAAGATATTGG | NNNTCATAGCTCCACYNGAATGWGCTARAATACC |
| NNTCGATACAGGTCAACAAATCATAAAGATATTGG | NNNNTCGATGCGCCACYNGAATGWGCTARAATACC |
| NNNTCTCATCAGGTCAACAAATCATAAAGATATTGG | NNTCTAGTGACCACYNGAATGWGCTARAATACC |
| NNNNTCTGCACTGGTCAACAAATCATAAAGATATTGG | NNNTGATGAGACCACYNGAATGWGCTARAATACC |
| NNTGACTAGCGGTCAACAAATCATAAAGATATTGG | NNNNTGTAGACTCCACYNGAATGWGCTARAATACC |
| NNNTGTGATGTGGTCAACAAATCATAAAGATATTGG | NNTGTGTGAGCCACYNGAATGWGCTARAATACC |

**Table S3**. Morphospecies recovered in Steele et al. (2018).

| Family | Species | strain |
| --- | --- | --- |
| Centropyxididae | *Centropyxis aculeata* | aculeata |
| Centropyxididae | *Centropyxis aculeata* | discoides |
| Centropyxididae | *Centropyxis constricta* | aerophila |
| Centropyxididae | *Centropyxis constricta* | spinosa |
| Netzeliidae | *Cyclopyxis kahli* |  |
| Difflugidae | *Difflugia amphora* |  |
| Difflugidae | *Difflugia bidens* |  |
| Difflugidae | *Difflugia elegans* |  |
| Difflugidae | *Difflugia fragosa* |  |
| Difflugidae | *Difflugia glans* | glans |
| Difflugidae | *Difflugia glans* | magna |
| Difflugidae | *Difflugia glans* | distenda |
| Difflugidae | *Difflugia globulosa* |  |
| Difflugidae | *Difflugia oblonga* | bryophila |
| Difflugidae | *Difflugia oblonga* | lanceolata |
| Difflugidae | *Difflugia oblonga* | linearis |
| Difflugidae | *Difflugia oblonga* | oblonga |
| Difflugidae | *Difflugia oblonga* | spinosa |
| Difflugidae | *Difflugia oblonga* | tenuis |
| Difflugidae | *Difflugia protaeiformis* | acuminata |
| Difflugidae | *Difflugia urceolata* | urceolata |
| Difflugidae | *Difflugia urens* |  |
| Netzeliidae | *Mediolus corona* |  |
| Difflugidae | *Lagenodifflugia vas* |  |
| Difflugidae | *Pontigulasia compressa* |  |
| Netzeliidae | *Cucurbitella tricuspis* |  |
| Difflugidae | *Lesquereusia spiralis* |  |

Table S4. Number of Illumina reads per sample through the dada2 pipeline.

| **Sample** | **input** | **filtered** | **denoisedF** | **denoisedR** | **merged** | **nonchim** | **%** |
| --- | --- | --- | --- | --- | --- | --- | --- |
| **Freshwater lake sediment** | | | | | | | |
| Q1 | 17584 | 15133 | 14884 | 14919 | 14308 | 14150 | 80,47 |
| Q2 | 9752 | 8508 | 8364 | 8358 | 7955 | 7790 | 79,88 |
| Q3 | 20645 | 18189 | 17675 | 17751 | 16821 | 16643 | 80,62 |
| ORO2 | 14824 | 11879 | 11617 | 11547 | 11223 | 11223 | 75,71 |
| FRO2 | 34378 | 30398 | 29739 | 29672 | 28036 | 27838 | 80,98 |
| **Estuary sediment** | | | | | | | |
| 13 | 34071 | 33420 | 33300 | 33247 | 32445 | 32130 | 94,3 |
| 35 | 32611 | 29312 | 28968 | 28751 | 27528 | 26156 | 80,21 |
| 48 | 26476 | 25817 | 25665 | 25540 | 25279 | 25227 | 95,28 |
| 78 | 54524 | 20053 | 19797 | 19597 | 18872 | 18439 | 33,82 |
| 79 | 28341 | 27209 | 27074 | 26932 | 26704 | 26672 | 94,11 |

Table S5. Number of reads per taxonomic assignation in each sample.

| **Freshwater lake sediment** | | | | | |
| --- | --- | --- | --- | --- | --- |
|  | **FRO2** | **ORO2** | **Q1** | **Q2** | **Q3** |
| Arcellinida | 20377 | 10177 | 6850 | 5237 | 12700 |
| Lobosa | 6941 | 853 | 5843 | 1826 | 3575 |
| “Other” | 208 | 16 | 12 | 0 | 8 |
| Oomycetes | 126 | 53 | 513 | 0 | 58 |
| SAR others | 39 | 62 | 858 | 679 | 175 |
| Total reads | 27691 | 11161 | 14076 | 7742 | 16516 |
| **Arcellinida percentage** | 0,735870 | 0,911835 | 0,486643 | 0,676440 | 0,768951 |
| **Estuary sediment** | | | | | |
|  | **79** | **78** | **48** | **35** | **13** |
| Lobosa | 25328 | 1438 | 24387 | 6777 | 25153 |
| Oomycetes | 941 | 4247 | 752 | 18036 | 6767 |
| Arcellinida | 274 | 15 | 10 | 11 | 36 |
| SAR others | 32 | 12575 | 46 | 1141 | 11 |
| “Other” | 0 | 26 | 0 | 0 | 0 |
| Total reads | 26575 | 18301 | 25195 | 25965 | 31967 |
| **Arcellinida percentage** | 0,010310 | 0,000819 | 0,000396 | 0,000423 | 0,001126 |
